# The effect of Rosovitine on mPSM explants: a real time analysis

**DOI:** 10.1101/789446

**Authors:** Lucas J. Morales Moya, Charlotte S. L. Bailey, J. Kim Dale, Philip J. Murray

## Abstract

Previously we showed, using fixed tissue techniques, that treatment of chick embryos with a family of pharmacological inhibitors yields increased levels of NICD, an increased NICD half life and longer segments (Wiederman et al., 2015). Here we measure the effect of one of the pharmacological perturbations (Roscovtine) using a real time reporter of the somitogenesis clock. After processing the reporter signal using empirical mode decomposition, we measure the oscillator period in mPSM explants and find, in agreement with the previous study, that the period of the segmentation clock increases upon Roscovitine treatment. However, we also make the novel discovery that the differentiation rate of the mPSM tissue also increases upon Roscovitine treatment. Returning to the previous study, we find that the measured increases in somite size and oscillator period are only consistent with the clock and wavefront model if the wavefront velocity also increased.

## 1 Introduction

During somitogenesis pairs of somites form periodically from the presomitic mesoderm, one on either side of the neural tube (Tam, 1981). The periodicity of somite formation is approximately constant for most of somitogenesis, taking approximately two hours in mice (?Lauschke et al., 2013), half an hour in zebrafish and an hour and a half in chick embryos. Whilst the number of somites, their size and the rate of formation is species dependent (Gomez et al., 2008), variation within individuals of the same species is less than 5% (Cooke, 1998).

Underlying periodic segment formation is an oscillatory pattern of gene expression that was first observed in the PSM of chick embryos (Palmeirim et al., 1997) but has since been identified in numerous vertebrate species e.g. chick (McGrew et al., 1998; Dubrulle et al., 2001; Dale et al., 2003), mouse (Bessho et al., 2001; Lewis, 2003; Morales et al., 2002; Bessho et al., 2003; Dequéant et al., 2006) and zebrafish (Holley and Takeda, 2002; Oates and Ho, 2002; Krol et al., 2011). Genes encoding the Hairy transcriptional repressor proteins that belong to the bHLH family of gene repressors, which are targets of the Notch pathway, are common to all studied vertebrate segmentation clock systems. However in the mouse and chick embryos components of the Fgf and Wnt signalling pathways have also been found to oscillate.

Notch is a juxtacrine cell-cell signalling pathway that regulates the transcription of hairy genes. Upon binding of Notch with Delta or Jagged of a neighbouring cell, Notch is cleaved and its intracellular domain, NICD, is transported into the nucleus where it is thought to form a transcriptional activator complex that activates the transcription of target genes (Fortini, 2009, see Figure 1). As cellular oscillators are subject to molecular noise, which disrupts synchrony between cells, it has been shown that one of the roles of Notch signalling is to synchronise oscillations in the PSM (Maroto et al., 2005; Masamizu et al., 2006; Riedel-Kruse et al., 2007; Ferjentsik et al., 2009; Özbudak and Lewis, 2008; Liao et al., 2016; Hubaud et al., 2017).

**Figure 1:**
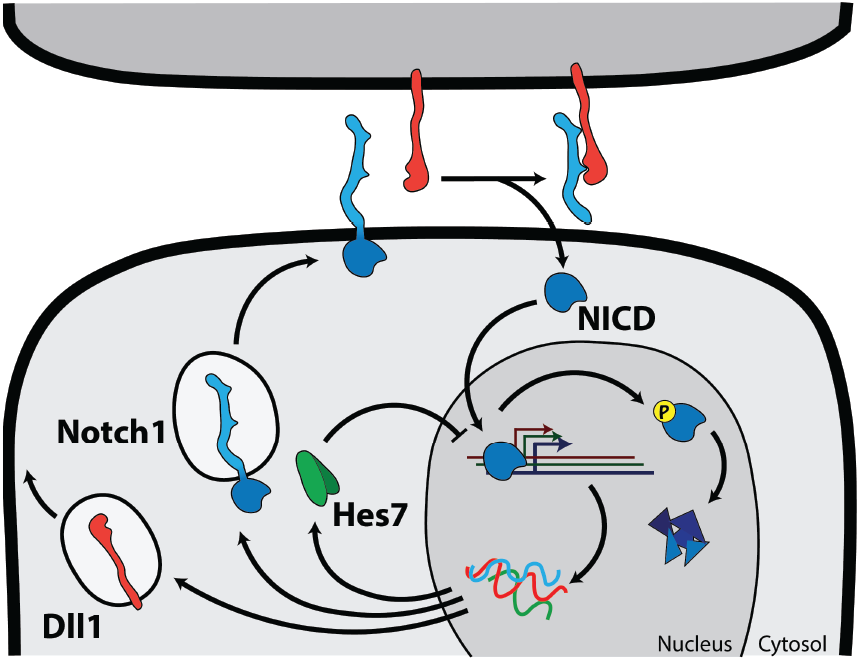
A schematic illustration of the role of Notch signalling in the mouse segmentation clock.

In a previous study in eLife Wiedermann et al. (2015) showed that pharmacological perturbations applied to PSM tissue had the following phenotype: longer NICD half life, higher levels of NICD and delayed formation of larger somites. Moreover, by reducing the production of NICD via treatment with a gamma secretase inhibitor, NICD levels and clock delay phenotype were rescued. Recent work in cell lines has demonstrated that the same pharmacological perturbations reduce the phosphorylation of NICD and hence its interaction with FBox7 which normally precedes NICD proteosomal degradation (Carrieri et al., 2019; Murray et al., 2019). The critical phosphosites required for Fbox7 interaction have been shown to be targets of the cyclin dependent kinases CDK1 and CDK2. These results suggest that CDK1 and CDK2 phosphorylation of NICD results in a decrease in its stability. This hypothesis was supported by experiments in which exposure of PSM explants to CDK specific inhibitors had the same phenotype as Roscovitine (i.e. higher levels of NICD and the delayed formation of larger somites).

The relatively recent development of of both real time reporters for clock gene expression and novel culture systems has enabled exciting phenomena to be uncovered (Delaune et al., 2012; Webb et al., 2016; Hubaud et al., 2017; Kageyama et al., 2015; Sonnen et al., 2018; Tsiairis and Aulehla, 2016; Lauschke et al., 2013; Soroldoni et al., 2014; Sonnen et al., 2018). Lauschke et al. (2013) showed that many of the features of segmentation clock dynamics that are observed *in vivo* (e.g. the emergence of waves of gene expression, segment formation, scaling phenomena) emerge in *ex vivo* mPSM tail explants.

Numerous methods have been used to process real time reporter signal so as to yield biologically meaningful inference (Lauschke et al., 2013; Webb et al., 2016; Hubaud et al., 2017; Tsiairis and Aulehla, 2016; Soroldoni et al., 2014; Sonnen et al., 2018; Matsumiya et al., 2018; Delaune et al., 2012). In recent work we proposed a method based on empirical mode decomposition that enable us to detrend and denoise a given signal in a systematic manner before applying a Hilbert transform in order to infer instantaneous phase (Morales et al., 2019). In order to characterise the effect of the pharmacological inhibitors used by Wiedermann et al. (2015) in a real time system, in this study we perform a set of experiments in which *ex vivo* mPSM explants (Lauschke et al., 2013) are treated with Roscovitine and compared with control samples. EMD-based methods are used to infer oscillator phase and a number of metrics are employed to quantify the effect of Roscovitine treatment.

## 2 Methods

### 2.1 Experimental

#### 2.1.1 Mouse line

The *LuVeLu* mouse (*Mus musculus*) (a gift from O. Pourquie) expresses *Venus-YFP* under the control of a 2.1kb fragment of the *Lunatic Fringe* (*Lfng*) promoter. The mRNA contains the 3’UTR of *Lfng* and the protein is fused to a PEST domain, to destabilise both the RNA and the protein and ensure clear oscillations (see Aulehla et al. (2008) for further details).

#### 2.1.2 Mating procedure

Mouse E10.5 embryos were generated and the line was maintained by crossing *LuVeLu* males to stock CD1 females, as the *LuVeLu* construct is lethal in homozygotes.

#### 2.1.3 Mouse genotyping

To renew the mouse line, timed matings were performed and the litters genotyped to ensure the *LuVeLu* construct is still present. A diagnostic PCR was performed on total DNA from an ear biopsy tissue of individual animals. DNA was extracted by incubation in microLYSIS-Plus buffer (*Thistle Scientific*) through the following PCR protocol: 65°C for 15 minutes, 96°C for 2 minutes, 65°C for 4 minutes, 96°C for 1 minute, 65°C for 1 minute, 96°C for 30 seconds and 8°C until stopped.

The PCR mix was generated by adding 2*µ*l of the lysed solution to a solution containing 1.25*µ*l GoTaq™Flexy polymerase (*Promega*), 0.31 mM dNTPs (*Promega*), 1.25 mM MgCl_2_, 1 GoTaq™Flexi PCR buffer (*Promega*) and 20 pmol of each of the following four primers: (i) Ala1 (Forward), 5’-tgctgctgcccgacaaccact-3’; (ii) Ala3 (Reverse), 5-tgaagaacacgactgcccagc-3; (iii) IMR0015, 5’-caaatgttgcttgtctggtg-3; (iv) IMR0016, 5-gtcagtcgagtgcacagttt-3. Distilled water was added to the solution to reach a final volume of 20*µ*l. The following PCR protocol was used: 94°C for 2 minutes; 35 cycles of [92°C for 45 seconds, 59°C for 40 seconds, 72°C for 40 seconds]; 75°C for 5 minutes; and 4°C until stopped.

PCR samples were analysed through electrophoresis, by loading 5*µ*l of each PCR product onto a 1% agarose gel with 1:10,000 Gel Red™(*Biotium/VWR*®) and run for 20 minutes at 100 volts. The result was visualised using an UV light box. Wild type CD1 presented a single fragment of 200 bp whilst *LuVeLu*^+^*/*− presented this fragment together with a 461 bp fragment.

### 2.1.4 ex vivo culture system

A 35 mm FluoroDish (*World Precission Instruments*™) with coverglass bottom was coated with a 50 *µ*g/ml fibronectin (*Sigma*) in a 100 mM sodium chloride (NaCl) solution made with double distilled water (ddH_2_O) before the dissection. The dish was incubated in the solution either for 4 hours at room temperature or overnight at 4 °C with agitation. The solution was later removed and the dish left until it was completely dried, around 30 minutes.

The dish was washed in tail bud culture medium of DMEM/F12 with no phenol red (*Gibco/Life Technologies*™) with 0.5 mM glucose (*Sigma*), 2mM L-glutamine (*Gibco/Life Technologies*™), 1% bovine serum albumin (BSA) (*Sigma*), penicillin/ streptomycin(*Gibco/Life Technologies*™) for 30 minutes at room temperature.

Embryos were harvested at E10.5 from timed matings. Individual embryos were taken from the uterine horn in sterile PBS using forceps and transferred to dissection media. To identify *luVeLu* positive embroys, tails were cut and transferred to a pre-warmed tail bud dissection media (tail bud culture media + 10 mM HEPES (*Sigma*) in a multi-well dish.

After *LuVeLu* tail identification, the tail bud was isolated from each tail posterior to the neuropore and transferred to an imaging disk, with the cut facing downwards, towards the fibronectin-coated surface.

Explants were incubated at 38.5°C, 5% CO_2_ for 1 hour to allow the explant to adhere to the fibronectin-coated surface, before live imaging. Embryos were transferred to a confocal microscope, as described in Section 2.1.5, and imaged for 24 hours at 37°C, 5% CO_2_ and ambient O_2_ levels.

#### 2.1.5 Imaging

Tail bud explants were prepared as described above. The imaging dish was transferred to a 37°C heated stage with a heated chamber at 37°C with 5% CO_2_ and ambient O_2_ of a Zeiss 710 inverted confocal microscope. Explants were imaged using a EC Plan-Neoufluar 10x/0.30 dry objective (*Zeiss*, M27). The *LuVeLu* fluorophore was excited by using a 514nm Argon laser.

Samples were scanned bi-directionally, to increase acquisition speed, and averaged 8 times per line, with a spatial resolution of 1024×1024 pixels and a temporal resolution of 15 minutes. Three optical planes were acquired at 14*µ*m intervals and starting at the plane of the cut, and moving upwards.

#### 2.1.6 Drug assay

Embryos were harvested and cultured as described above. However, to allow for different conditions to be tested in the same disk, Multi-well silicone inserts (*Culture insets; iBidi*) were adhered to a 35 mm FluoroDish (*World Precission Instruments*™), prior to the fibronectin coating.

Roscovitine (*Calbiochem*), previous dissolved in DMSO at a concentration of 10*µ*M, was added to the tail bud culture media in a 1:1000 relationship and put aside.

Tail bud culture media was replaced after 16 hours of imaging with tail bud culture media that contained 0.1*µ*M Roscovitine tailbud media with or an equivalent volume of DMSO. The imaging was resumed until *t* = 24 hours after re-estabilishing the optical planes to compensate for perturbations in the *z*-axis after the media addition.

### 2.2 Phase reconstruction

Let *s*(x*, t*) represent a spatio-temporal oscillatory signal measured from a realtime reporter from an mPSM explant. The main steps used to generate the reconstructed phase profile (see Figure 2) are outlined below.

**Figure 2:**
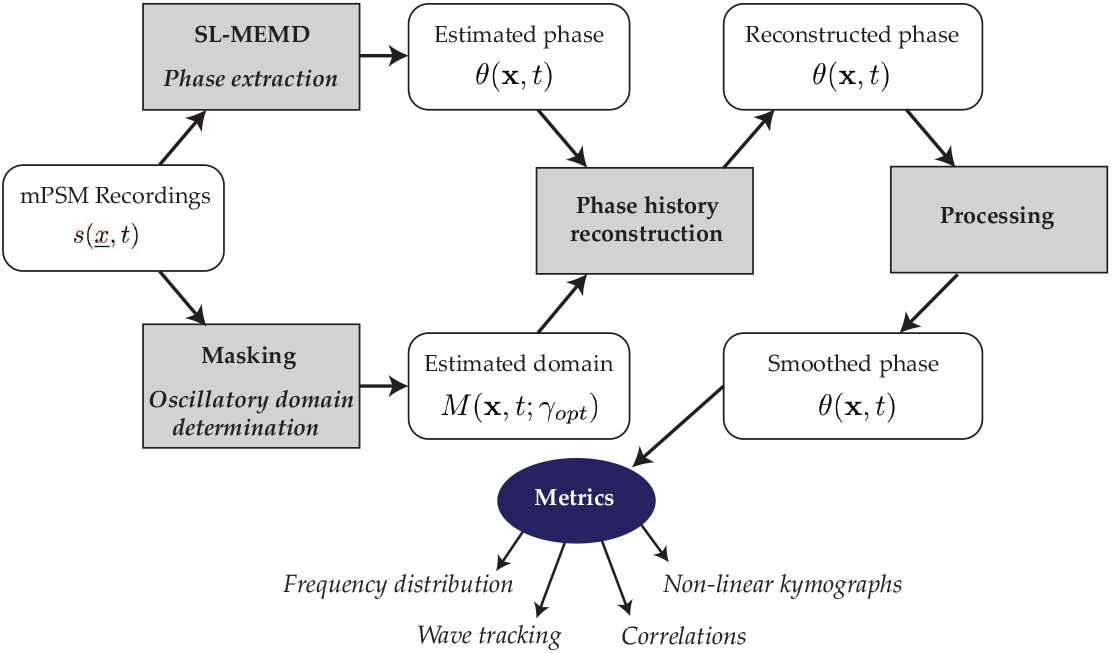
A schematic illustration of phase reconstruction.

Oscillator phase is reconstructed from real time reporter signal using SLMEMD (Morales et al., 2019). A thresholding algorithm was used to define the actively oscillating region, *M* (*<u>x</u>*, *t*), and phase is fixed on the segmenting boundary when signal is lost. Further details can be found in Morales et al. (2019).

### 2.3 Metric computation

#### 2.3.1 Wave tracking

A wave tracking algorithm was used to identify the emergence of each successive wave of gene expression. By measuring the time that elapses between successive waves the tissue scale period of the pattern was computed. Given that the unwrapped oscillator phase can be written in the form

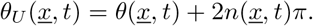

the number of elapsed cycles in the actively oscillating domain is given by

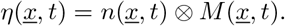

Defining the number of actively oscillating voxels at time *t* to be

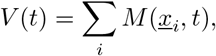

the fraction of voxels with *k* elapsed cycles is given by

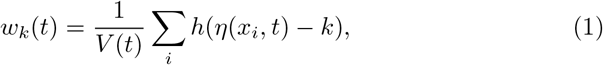

and *h*(.) is an indicator function. The time of the emergence of the k^th^ wave, 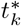, is defined to be

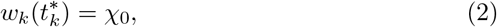

where *χ*_0_ is a constant. The tissue scale period is defined to be the time between the emergence of consecutive waves, i.e.

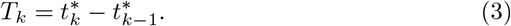

#### 2.3.2 Differentiation rate

Defining *V* (*t*) to be the number of actively oscillating voxels at time *t*, the averaged normalised differentiation rate is defined to be

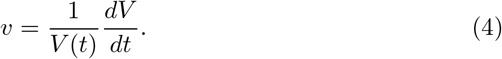

## 3 Results

To quantify the effect of Roscovitine treatment on clock gene expression, mPSM explants isolated from E10.5 LuVeLu embryos were treated with 20 *µ*M Roscovitine or a DMSO control at *t* = 15h, cultured on fibronectin coated glass slides (Section 2.1)) and imaged for up to 24 hours using an Zeiss 710 confocal microscope.

We found that mPSM samples showed qualitatively similar behaviour to that described in Morales et al. (2019) (see Appendix A). In the Roscovitine-treated samples, fluoresence intensity from the LuVeLu reporter was dynamic (see Figure 3 (a)), and, upon moving average subtraction, exhibited clear oscillatory waves (see Figure 3 (b)) on a dynamic spatial domain (see Figure 3 (c)). After using SLMEMD to reconstruct the oscillator phase for each mPSM explant (see Figure 3 (d) and Section 2.2), we identified, as previously reported, initially spatially homogeneous oscillations. Subsequently, oscillations were halted on the periphery of the mPSM explant and waves of gene expression were observed travelling from the centre to the periphery of the explant.

**Figure 3:**
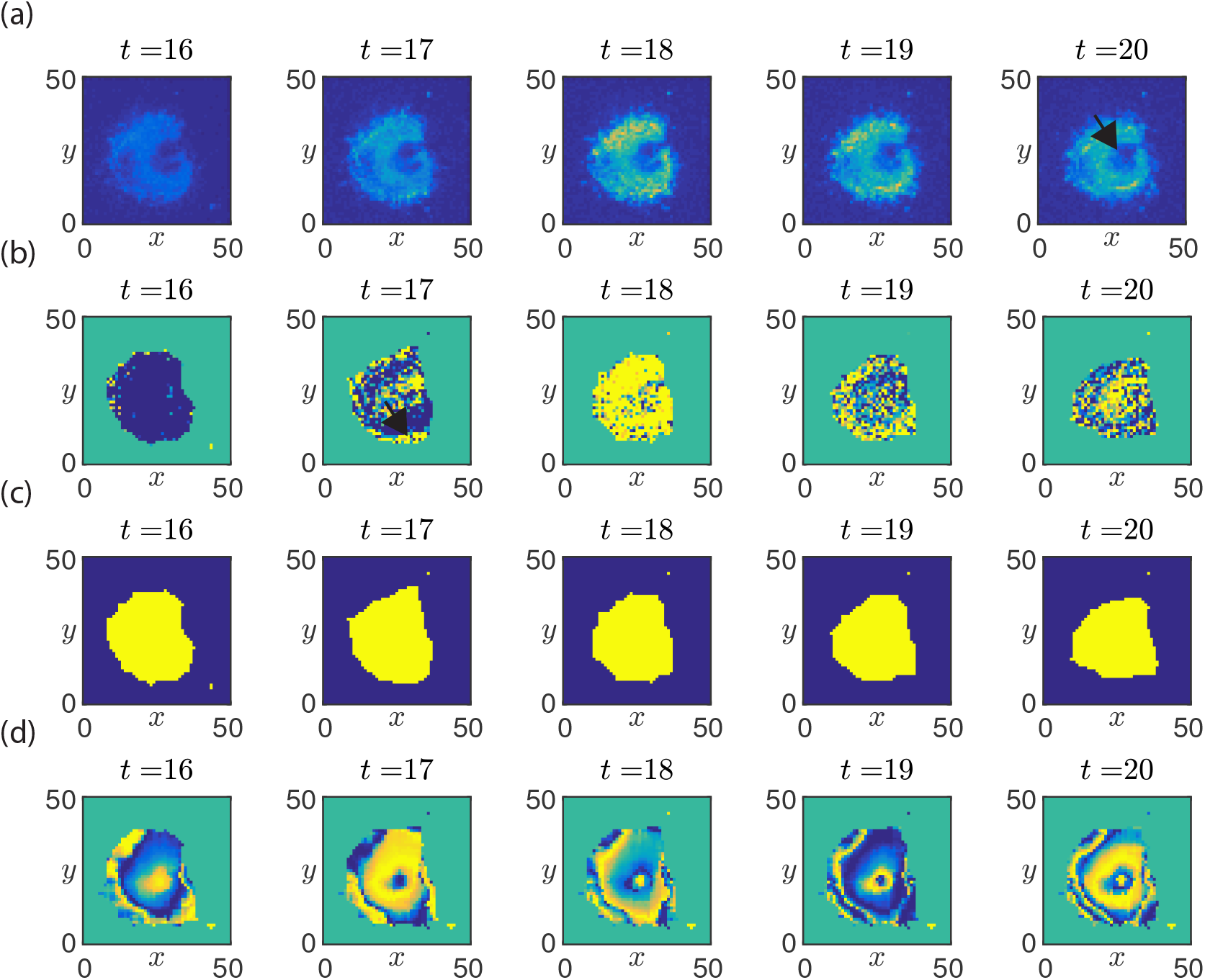
Spatiotemporal dynamics in a Roscovitine treated mPSM explant. (a) Snapshots of the fluorescent signal from the LuVeLu reporter. (b) Snapshots of the fluorescent signal after application of a moving average filter. (c) Snapshots of the mask that defined the extent of the actively signalling domain. (d) Snapshots of the sine of the reconstructed oscillator phase, 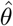, that was computed using the method outlined in Figure 2. Unit length - 10*µ*m. Black arrows represent direction of propagation. Yellow - high, Blue-low.

Using the wave tracking algorithm (see Section 2.3.1) we identified the time of emergence of successive oscillatory waves in each of the mPSM samples (see 4 (a) and (b)) and, after computing the time difference between the emergence of successive waves, identified the tissue scale frequency of each mPSM sample (see 4 (c)). To characterise the differentiation rate of the tissue, we measured the area of the actively oscillating region (see Section 2.3.2) as a function of time (see Figure 4 (d)).

**Figure 4:**
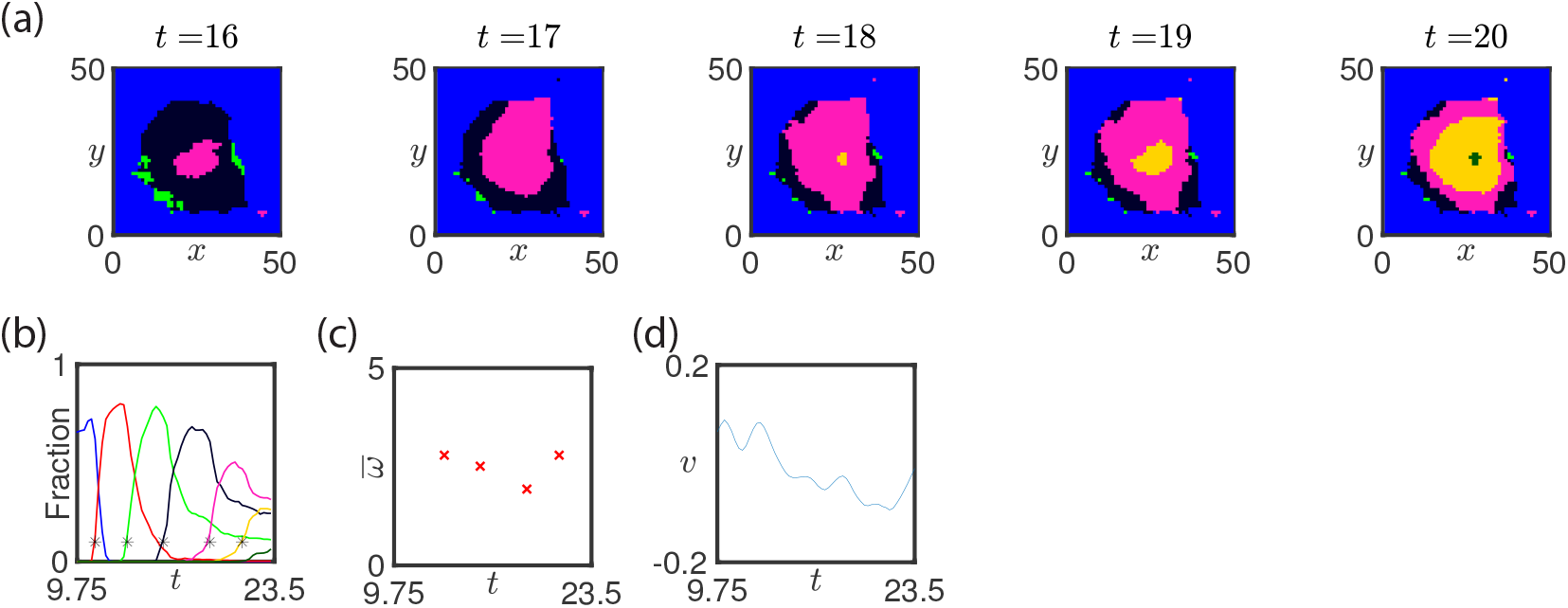
Quantitative analysis of a Roscovitine treated mPSM explant. (a) Snapshots of the emergence of successive waves (labelled with different colours). (b) The area spanned by each wave is plotted against time (colours represent wave numbers plotted in (a), markers represent initiation of new wave). (c) The inter-wave period is plotted against time (see equation (3)). (d) The average differentiation rate is plotted against time (equation (4)).

To quantify the relative effect of Roscovitine treatment, we performed a statistical analysis of the tissue scale frequency and differentiation rate in both control and Roscovitine-treated groups. Using the wave tracking algorithm we found that the Roscovitine treated group had a reduced tissue scale frequency relative to the DMSO control group after Roscovitine was added (*t* = 16*h*) but that the effect was transient. Using the differentiate rate tracking algorithm, we found that the Roscovitine treatment group had an increased differentiation rate relative to the control group (see Figure 5 (b)). These results show, for the first time in a real time reporter system, that Roscovitine treatment does indeed reduce the tissue scale frequency of oscillation in mPSM explants. Moreover, we have made the novel finding that Roscovitine treatment also increases the differentiation rate of the tissue.

**Figure 5:**
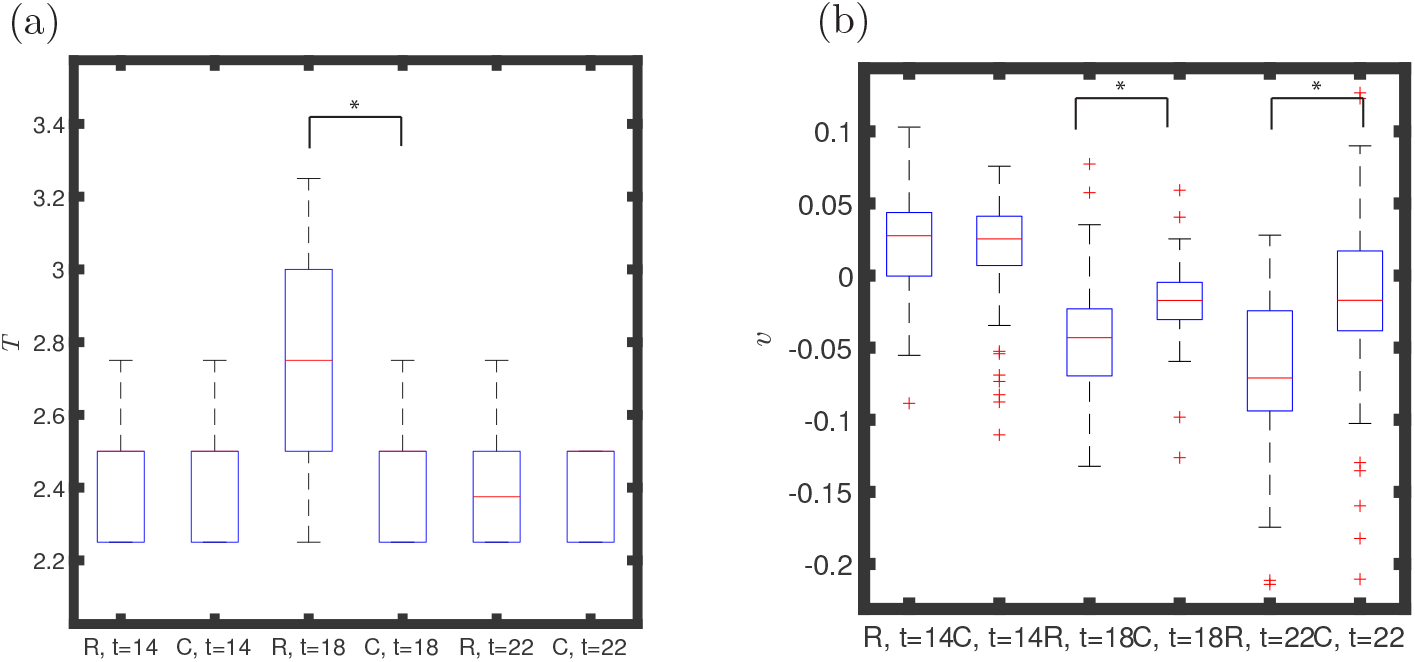
Quantitative analysis of mPSM explants upon Roscovitine treatment. (a) Box plots of tissue scale period measurements, *T*, are plotted at four hour intervals for Roscovitine (R) and DMSO treated (C) mPSM samples. (b) Box plots of the differentiation rate, *v*, plotted at four hour intervals for Roscovitine (R) and DMSO (C)treated samples. Roscovitine (*n* = 6). DMSO (*n* = 6).

## 4 Conclusions

During segmentation of the vertebrate embryo, pairs of somites form at regular intervals in time that are regulated a by a coupled molecular oscillator known as the somitogenesis clock.

In Wiedermann et al. (2015) we found that treatment of chick or mouse PSM with a family of pharmacological inhibitors results in a consistent phenotype: higher levels of NICD, an increased NICD half life, larger segments and an increased oscillation period. Using mathematical modelling we identified that reducing the production rate of NICD in the system would balance the increased stability of NICD so as to rescue the increased period phenotype. When we experimentally tested that prediction by simultaneously treating chick embryos with both Roscovitine and the gamma secretase inhibitor LY-374973, we observed partial rescue of the phenotype as predicted by the model.

In the Wiedermann et al. (2015) study, PSM halves from the same embryo were cultured in either DMSO or Roscovitine, then fixed and analysed by *in situ* for *Lnfg* RNA expression and the relative effect of the perturbation was inferred via comparison of clock gene expression patterns in the contralateral PSM halves. Moreover, the clock period was inferred in a given treatment condition using a ‘fix and culture’ experiment whereby one PSM half is fixed at a given time and the contralateral half is cultured until the phase pattern has progressed through a cycle. The inference from such experiments is limited by the fact that the oscillations cannot be measured directly in a single tissue sample. Moreover, other features of the spatio-temporal patterns, such as the length scale and position of the phase gradient, cannot be measured. Here we address these limitations by measuring the effect of one of the pharmacological treatments considered by Wiedermann et al. (2015) (Roscovitine) in a real time reporter system.

To assay the effect of Roscovitine treatment in real time, mPSM explants harvested from E10.5 LuVeLu embryos were separated into two groups, subjected to either DMSO or Roscovitine treatment and the fluorescent signal from the LuVeLu reporter was recorded. After an initial attempt at a naive peak-to-peak method measurement to infer the oscillation period that could not detect a statistically significant Roscovitine treatment effect, we then developed and applied the EMD-derived method to reconstruct the phase profile in space and time and computed a number of metrics that allowed biologically meaningfully quantities to be measured. When we applied the developed methodology to control samples, we detected an average oscillation rate that was consistent with previous methods (Lauschke et al., 2013). Consistent with the observations of Roscovitine treatment by Wiedermann et al. (2015) in chick and mouse embryos, we observed an increased period of the segmentation clock relative to DMSO in mPSM explants. Additionally, we made the novel finding that the differentiation rate of mPSM tissue is also increased relative to control explants following exposure to Roscovitine.

It is notable that the relative change in period measured in mouse mPSM explants (10% − 15%) is considerably smaller than that observed in chick embryos by Wiedermann et al. (2015) (∼ 33%). Whilst we do not not know whether this discrepancy is species specific or a function of the culture system, the fact that the effect of Roscovitine treatment is much smaller in the *ex vivo* mouse system goes some way to explaining why the effect was not detected using the naive approach.

Having identified an effect on the clock period and differentiation rate in mPSM explants, we returned to the Wiedermann et al. (2015) experiments to further explore whether the measured change in segment size could be predicted from the measured clock period using the clock and wavefront model. Given the measurements presented in Wiedermann et al. (2015), in the chick embryo we compute that the somite length in the +3 position changes by a factor of 1.5 whilst the period increases by a factor of 1.3. However, in the clock and wavefront model we expect to find that the control and Roscovitine treated segment lengths satisfy

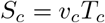

and

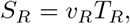

where *v*_*c*_ and *v*_*R*_ are the wavefront velocities and *T*_*c*_ and *T*_*R*_ are the clock periods, respectively. Hence in a situation where the wavefront speed is unaffected by Roscovitine treatment, we expect to find that the relative change in clock period matches the relative change in somite length, i.e.

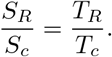

As in Wiedermann et al. (2015) the measured change in somite length is greater than the measured change in clock period, the clock and wavefront model is only consistent with the experimental data if the wavefront speed also increases. The computed results suggest the increase in somite size to be around 1.8 times larger when perturbed using Roscovitine. It remains to be seen if and how Roscovitine effects on differentiation rate are mediated by changes in stability levels of NICD.

In our hands we found a surprisingly large variability in the phase patterns in mPSM explants. After reconstructing phase profiles it is clear that the pattern geometry and boundary conditions have a pronounced effect on the geometry of the phase patterns observed within the tissue. These observations will be explored using mathematical modelling in a future study.

## A Phase reconstruction for a representative selection of mPSM samples

Spatio-temporal dynamics of DMSO-treated samples are presented in Figures 6–8.

**Figure 6:**
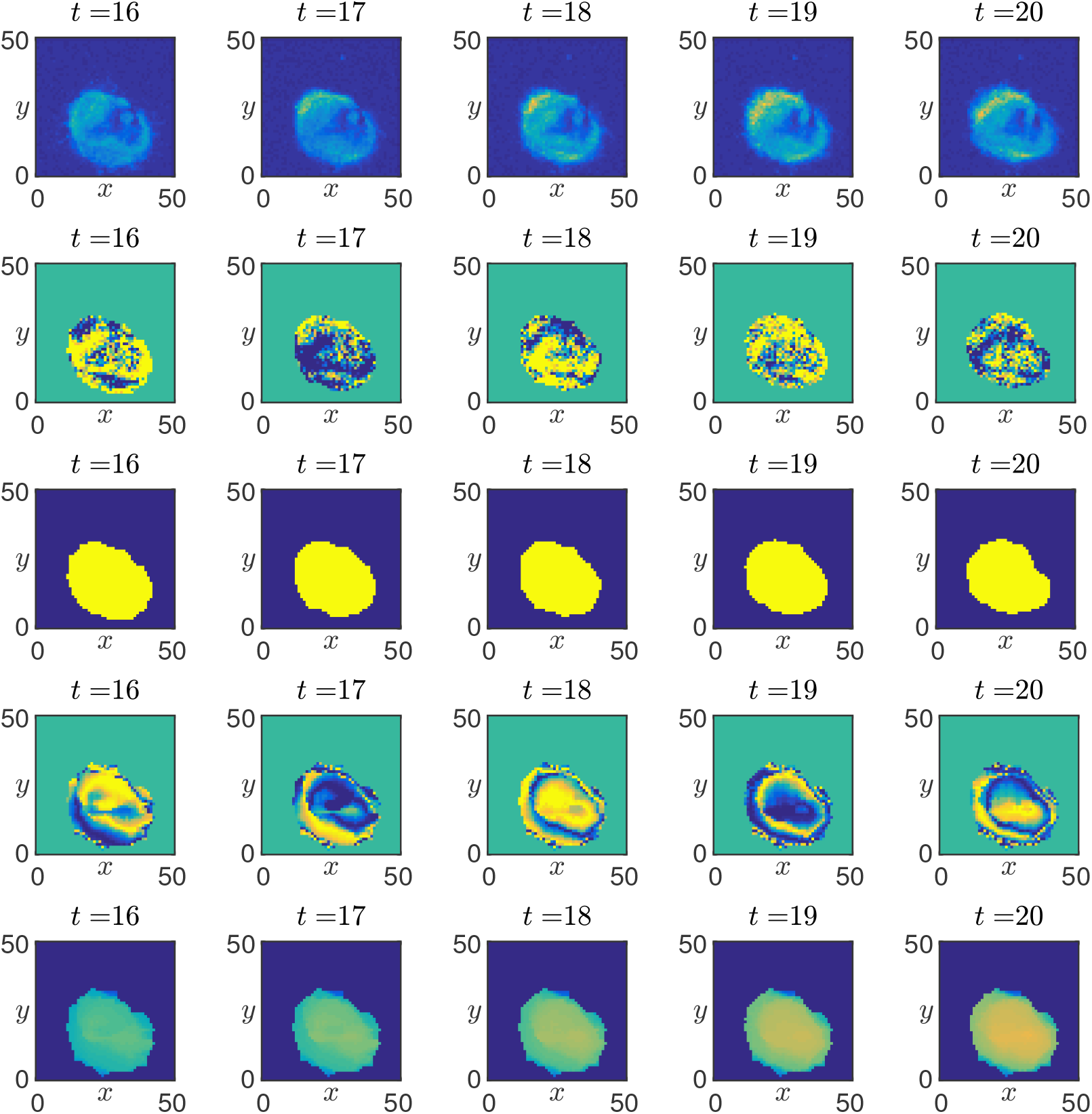
Quantitative analysis of oscillatory dynamics in mPSM explants. (a) Snapshots of the fluorescent signal from the LuVeLu reporter. (b) Snapshots of the fluorescent signal after application of a moving average filter. (c) Snapshots of the mask that defined the extent of the actively signalling domain. (d) Snapshots of the sine of the reconstructed oscillator phase, 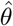, that was computed using the method outlined in Figure 2.

**Figure 7:**
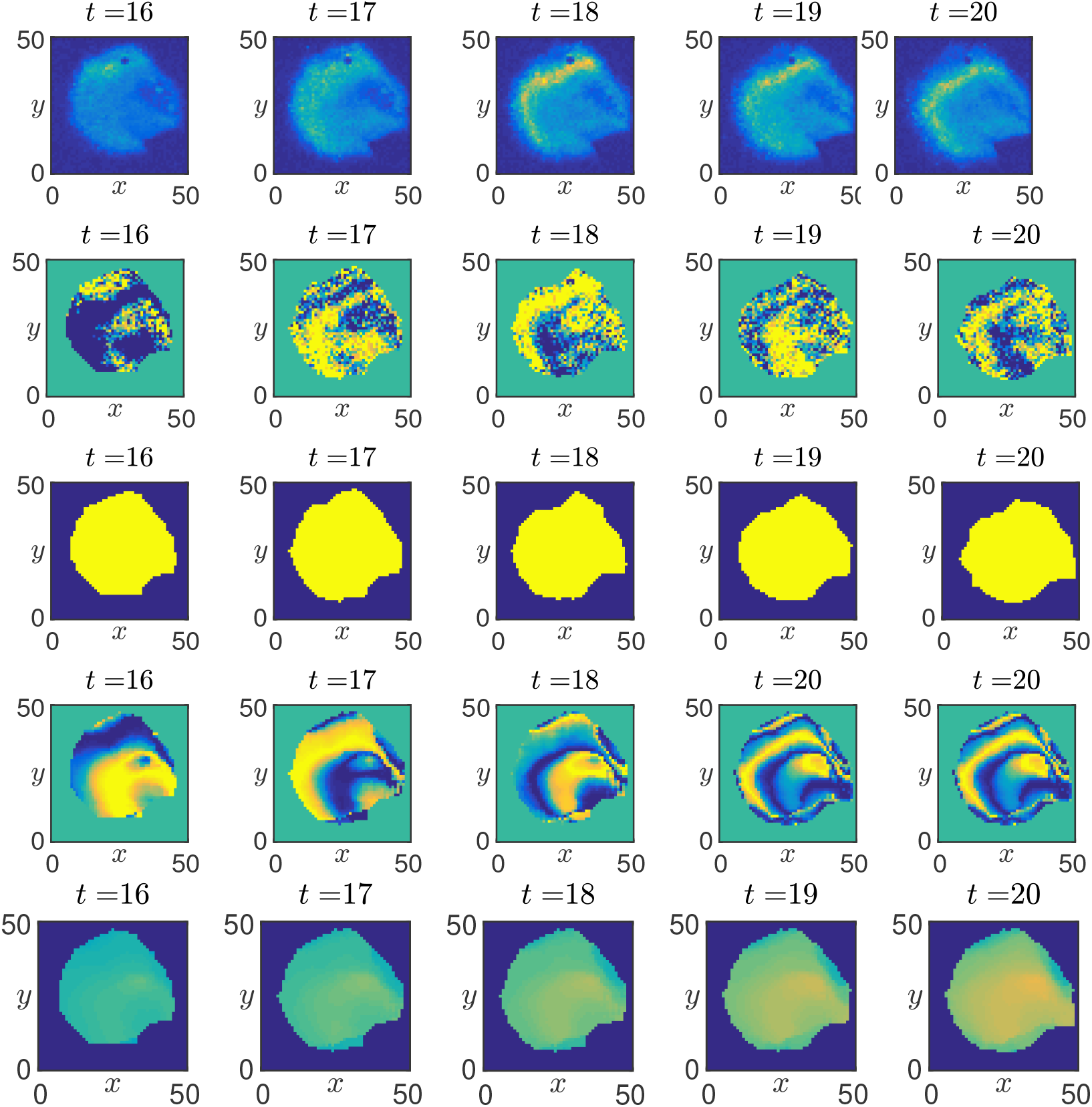
Quantitative analysis of oscillatory dynamics in mPSM explants. (a) Snapshots of the fluorescent signal from the LuVeLu reporter. (b) Snapshots of the fluorescent signal after application of a moving average filter. (c) Snapshots of the mask that defined the extent of the actively signalling domain. (d) Snapshots of the sine of the reconstructed oscillator phase, 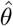, that was computed using the method outlined in Figure 2.

**Figure 8:**
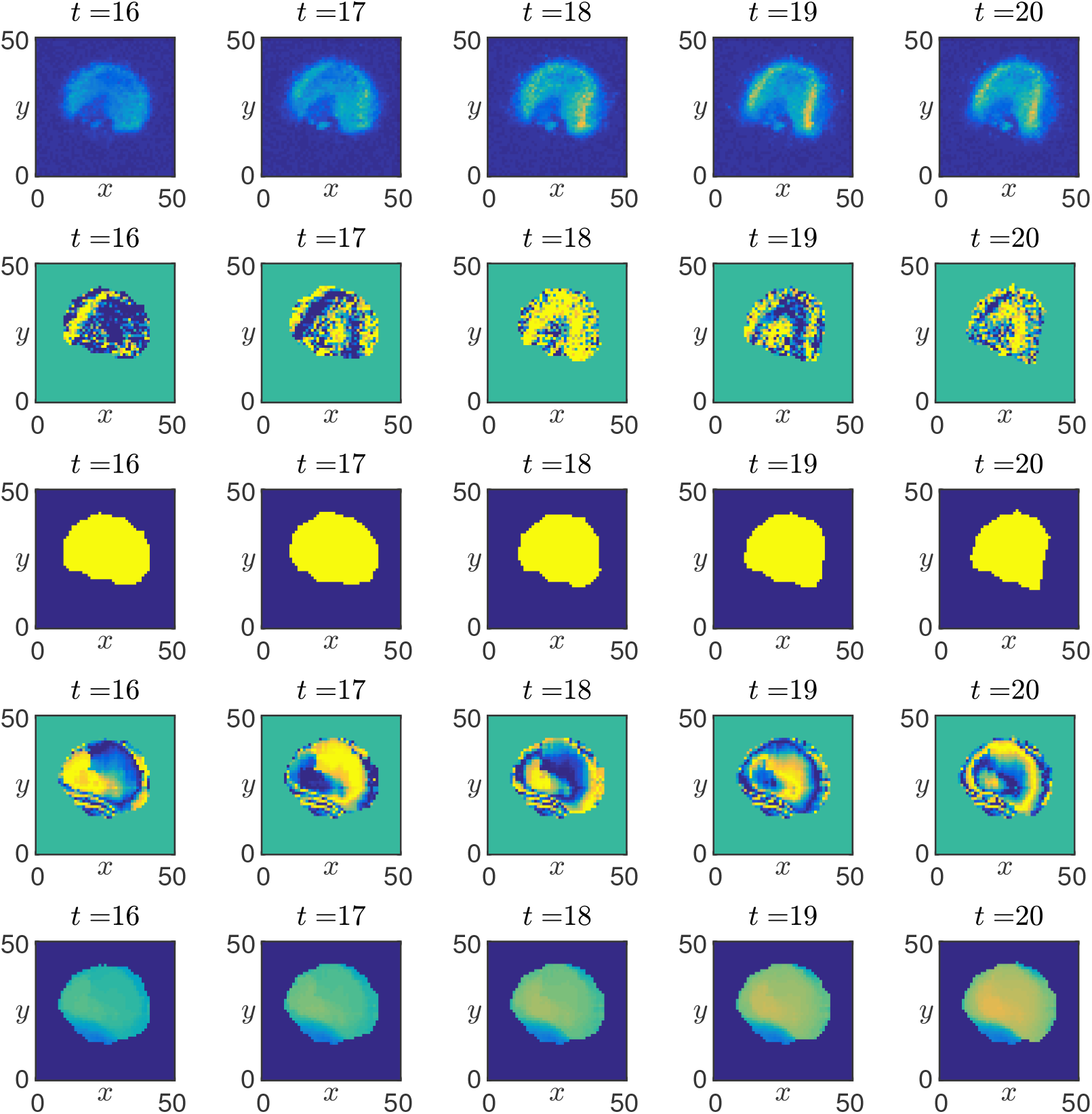
Quantitative analysis of oscillatory dynamics in mPSM explants. (a) Snapshots of the fluorescent signal from the LuVeLu reporter. (b) Snapshots of the fluorescent signal after application of a moving average filter. (c) Snapshots of the mask that defined the extent of the actively signalling domain. (d) Snapshots of the sine of the reconstructed oscillator phase, 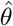, that was computed using the method outlined in Figure 2.

Spatio-temporal dynamics of Roscovitine-treated samples are presented in Figures 9 – 10.

**Figure 9:**
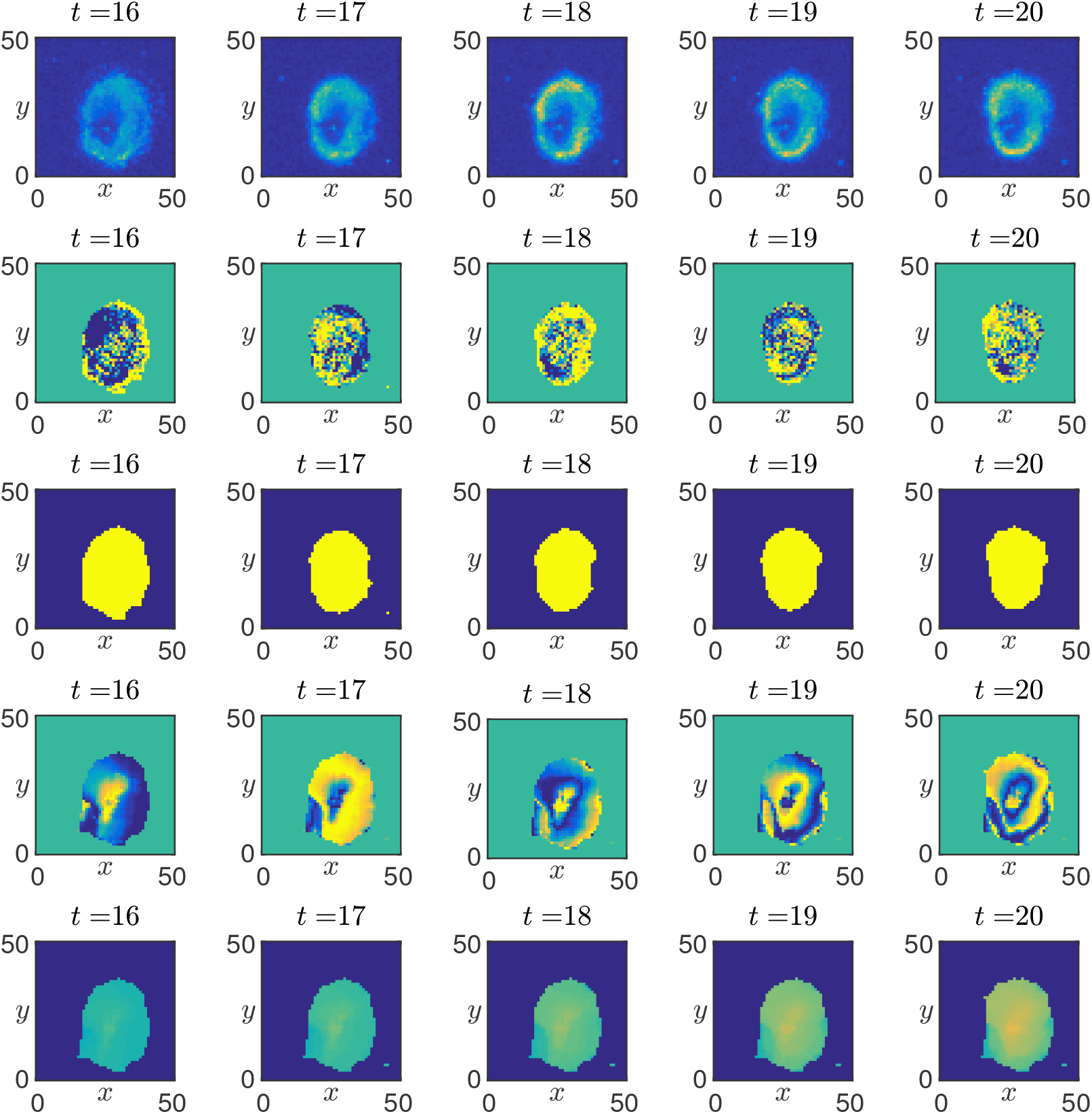
Quantitative analysis of oscillatory dynamics in mPSM explants. (a) Snapshots of the fluorescent signal from the LuVeLu reporter. (b) Snapshots of the fluorescent signal after application of a moving average filter. (c) Snapshots of the mask that defined the extent of the actively signalling domain. (d) Snapshots of the sine of the reconstructed oscillator phase, 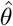, that was computed using the method outlined in Figure 2.

**Figure 10:**
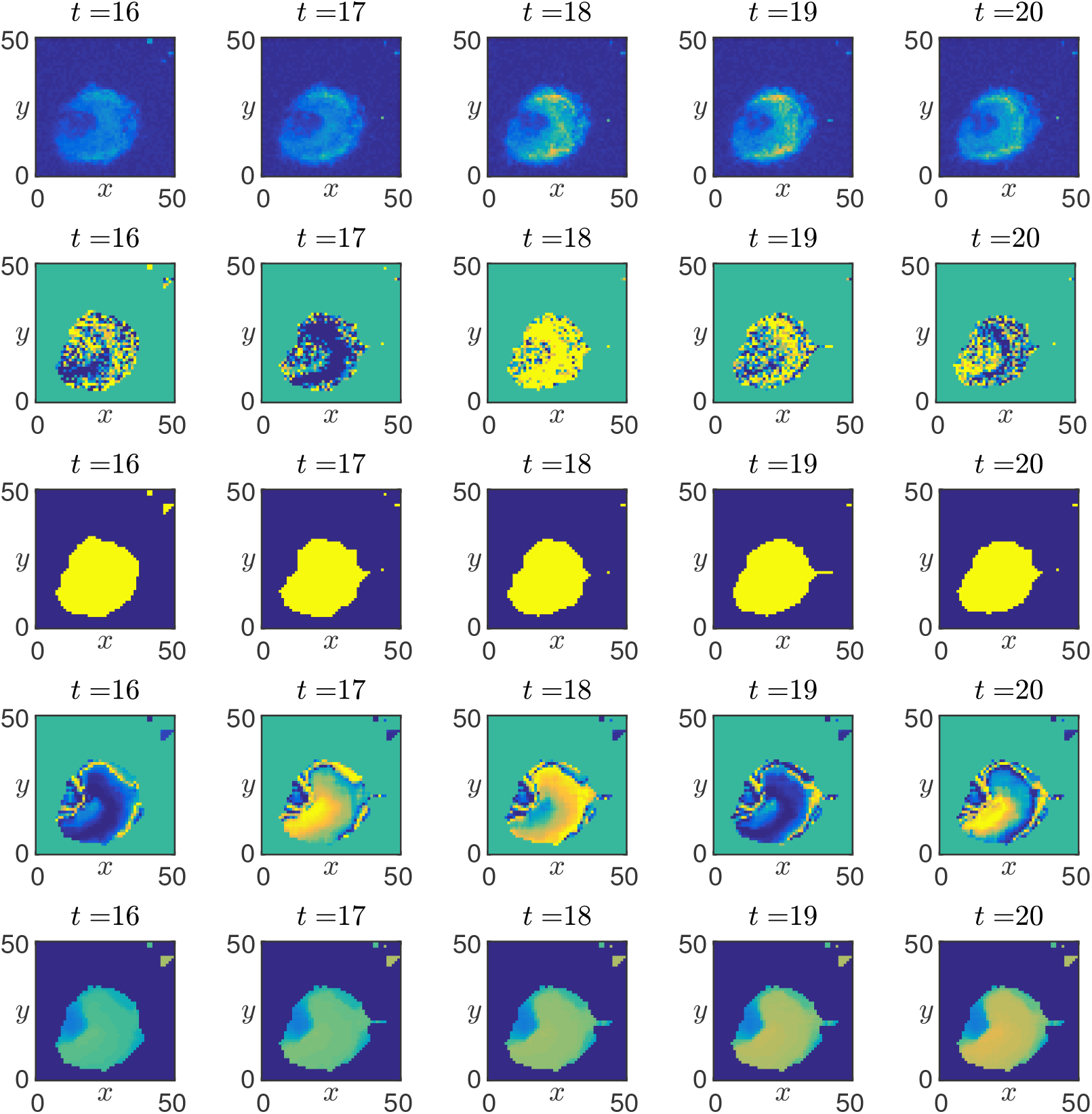
Quantitative analysis of oscillatory dynamics in mPSM explants. (a) Snapshots of the fluorescent signal from the LuVeLu reporter. (b) Snapshots of the fluorescent signal after application of a moving average filter. (c) Snapshots of the mask that defined the extent of the actively signalling domain. (d) Snapshots of the sine of the reconstructed oscillator phase, 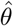, that was computed using the method outlined in Figure 2.

## References

Alexander Aulehla, Winfried Wiegraebe, Valerie Baubet, Matthias B Wahl, Chuxia Deng, Makoto Taketo, Mark Lewandoski, and Olivier Pourquié. A β-catenin gradient links the clock and wavefront systems in mouse embryo segmentation. Nature Cell Biology, 10(2):186, 2008.

Yasumasa Bessho, Goichi Miyoshi, Ryoichi Sakata, and Ryoichiro Kageyama. Hes7: a bHLH-type repressor gene regulated by Notch and expressed in the presomitic mesoderm. Genes to Cells, 6(2):175–185, 2001.

Yasumasa Bessho, Hiromi Hirata, Yoshito Masamizu, and Ryoichiro Kageyama. Periodic repression by the bHLH factor Hes7 is an essential mechanism for the somite segmentation clock. Genes & Development, 17(12):1451–1456, 2003.

Francesca A Carrieri, Philip J. Murray, Dimitrinka Ditsova, Margaret A Ferris, Paul Davies, and J Kim Dale. CDK1 and CDK2 regulate phosphorylation-dependent NICD1 turnover and the periodicity of the segmentation clock. EMBO reports, 2019.

Jonathan Cooke. A gene that resuscitates a theorysomitogenesis and a molecular oscillator. Trends in Genetics, 14(3):85–88, 1998.

J Kim Dale, Miguel Maroto, M-L Dequeant, Pascale Malapert, M McGrew, and Olivier Pourquie. Periodic Notch inhibition by lunatic fringe underlies the chick segmentation clock. Nature, 421(6920):275–278, 2003.

Emilie A Delaune, Paul François, Nathan P Shih, and Sharon L Amacher. Single-cell-resolution imaging of the impact of Notch signaling and mitosis on segmentation clock dynamics. Developmental Cell, 23(5):995–1005, 2012.

Mary-Lee Dequéant, Earl Glynn, Karin Gaudenz, Matthias Wahl, Jie Chen, Arcady Mushegian, and Olivier Pourquié. A complex oscillating network of signaling genes underlies the mouse segmentation clock. Science, 314(5805): 1595–1598, 2006.

Julien Dubrulle, Michael J McGrew, and Olivier Pourquié. FGF signaling controls somite boundary position and regulates segmentation clock control of spatiotemporal Hox gene activation. Cell, 106(2):219–232, 2001.

Zoltan Ferjentsik, Shinichi Hayashi, J Kim Dale, Yasumasa Bessho, An Herreman, Bart De Strooper, Gonzalo del Monte, Jose Luis de la Pompa, and Miguel Maroto. Notch is a critical component of the mouse somitogenesis oscillator and is essential for the formation of the somites. PLoS Genetics, 5 (9):e1000662, 2009.

Mark E Fortini. Notch signaling: the core pathway and its posttranslational regulation. Developmental Cell, 16(5):633–647, 2009.

Céline Gomez, Ertuğrul M Özbudak, Joshua Wunderlich, Diana Baumann, Julian Lewis, and Olivier Pourquié. Control of segment number in vertebrate embryos. Nature, 454(7202):335–339, 2008.

Scott A Holley and Hyroyuki Takeda. Catching a wave: the oscillator and wavefront that create the zebrafish somite. In Seminars in Cell & Developmental Biology, volume 13, pages 481–488, 2002.

Alexis Hubaud, Ido Regev, L Mahadevan, and Olivier Pourquie. Excitable dynamics and Yap-dependent mechanical cues drive the segmentation clock. Cell, 171(3):668–682, 2017.

Ryoichiro Kageyama, Hiromi Shimojo, and Itaru Imayoshi. Dynamic expression and roles of Hes factors in neural development. Cell and Tissue Research, 359 (1):125–133, 2015.

Aurélie J Krol, Daniela Roellig, Mary-Lee Dequéant, Olivier Tassy, Earl Glynn, Gaye Hattem, Arcady Mushegian, Andrew C Oates, and Olivier Pourquié. Evolutionary plasticity of segmentation clock networks. Development, 138 (13):2783–2792, 2011.

Volker M Lauschke, Charisios D Tsiairis, Paul François, and Alexander Aulehla. Scaling of embryonic patterning based on phase-gradient encoding. Nature, 493(7430):101, 2013.

Julian Lewis. Autoinhibition with transcriptional delay: a simple mechanism for the zebrafish somitogenesis oscillator. Current Biology, 13(16):1398–1408, 2003.

Bo-Kai Liao, David J Jörg, and Andrew C Oates. Faster embryonic segmentation through elevated delta-notch signalling. Nature Communications, 7, 2016.

Miguel Maroto, J Kim Dale, Mary-Lee Dequéant, Anne-Cecile Petit, and Olivier Pourquié. Synchronised cycling gene oscillations in presomitic mesoderm cells require cell-cell contact. International Journal of Developmental Biology, 49 (2-3):309–315, 2005.

Yoshito Masamizu, Toshiyuki Ohtsuka, Yoshiki Takashima, Hiroki Nagahara, Yoshiko Takenaka, Kenichi Yoshikawa, Hitoshi Okamura, and Ryoichiro Kageyama. Real-time imaging of the somite segmentation clock: revelation of unstable oscillators in the individual presomitic mesoderm cells. Proceedings of the National Academy of Sciences, 103(5):1313–1318, 2006.

Marina Matsumiya, Takehito Tomita, Kumiko Yoshioka-Kobayashi, Akihiro Isomura, and Ryoichiro Kageyama. ES cell-derived presomitic mesoderm-like tissues for analysis of synchronized oscillations in the segmentation clock. Development, 145(4), 2018.

Michael J Mcrew, J Kim Dale, Sandrine Fraboulet, and Olivier Pourquié. The lunatic fringe gene is a target of the molecular clock linked to somite segmentation in avian embryos. Current Biology, 8(17):979–982, 1998.

Aixa V Morales, Yuko Yasuda, and David Ish-Horowicz. Periodic lunatic fringe expression is controlled during segmentation by a cyclic transcriptional enhancer responsive to notch signaling. Developmental Cell, 3(1):63–74, 2002.

L. Morales, J. Kim Dale, and Philip J Murray. Reconstruction of the phase dynamics of the somitogenesis clock oscillator. BioArchiv (BIORXIV/2019/743724), 2019.

PJ Murray, FA Carrieri, and JK Dale. Cell cycle regulation of oscillations yields coupling of growth and form in a computational model of the presomitic mesoderm. Journal of Theoretical Biology, 2019.

Andrew C Oates and Robert K Ho. Hairy/E (spl)-related (Her) genes are central components of the segmentation oscillator and display redundancy with the Delta/Notch signaling pathway in the formation of anterior segmental boundaries in the zebrafish. Development, 129(12):2929–2946, 2002.

Ertuğrul M Özbudak and Julian Lewis. Notch signalling synchronizes the zebrafish segmentation clock but is not needed to create somite boundaries. PLoS Genetics, 4(2):e15, 2008.

Isabel Palmeirim, Domingos Henrique, David Ish-Horowicz, and Olivier Pourquié. Avian hairy gene expression identifies a molecular clock linked to vertebrate segmentation and somitogenesis. Cell, 91(5):639–648, 1997.

Ingmar H Riedel-Kruse, Claudia Müller, and Andrew C Oates. Synchrony dynamics during initiation, failure, and rescue of the segmentation clock. Science, 317(5846):1911–1915, 2007.

Katharina F Sonnen, Volker M Lauschke, Julia Uraji, Henning J Falk, Yvonne Petersen, Maja C Funk, Mathias Beaupeux, Paul François, Christoph A Merten, and Alexander Aulehla. Modulation of phase shift between Wnt and Notch signaling oscillations controls mesoderm segmentation. Cell, 172 (5):1079–1090, 2018.

Daniele Soroldoni, David J Jörg, Luis G Morelli, David L Richmond, Johannes Schindelin, Frank Jülicher, and Andrew C Oates. A Doppler effect in embryonic pattern formation. Science, 345(6193):222–225, 2014.

PPL Tam. The control of somitogenesis in mouse embryos. Development, 65: 103–128, 1981.

Charisios D Tsiairis and Alexander Aulehla. Self-organization of embryonic genetic oscillators into spatiotemporal wave patterns. Cell, 164(4):656–667, 2016.

Alexis B Webb, Iván M Lengyel, David J Jörg, Guillaume Valentin, Frank Jülicher, Luis G Morelli, and Andrew C Oates. Persistence, period and precision of autonomous cellular oscillators from the zebrafish segmentation clock. eLife, 5:e08438, 2016.

Guy Wiedermann, Robert Alexander Bone, Joana Clara Silva, Mia Bjorklund, Philip J Murray, and J Kim Dale. A balance of positive and negative regulators determines the pace of the segmentation clock. eLife, 4:e05842, 2015.

